## Supplemental Fig 1-5 for "Heparan Sulfate-dependent RAGE oligomerization is indispensable for pathophysiological functions of RAGE"

Supplemental Figure S1

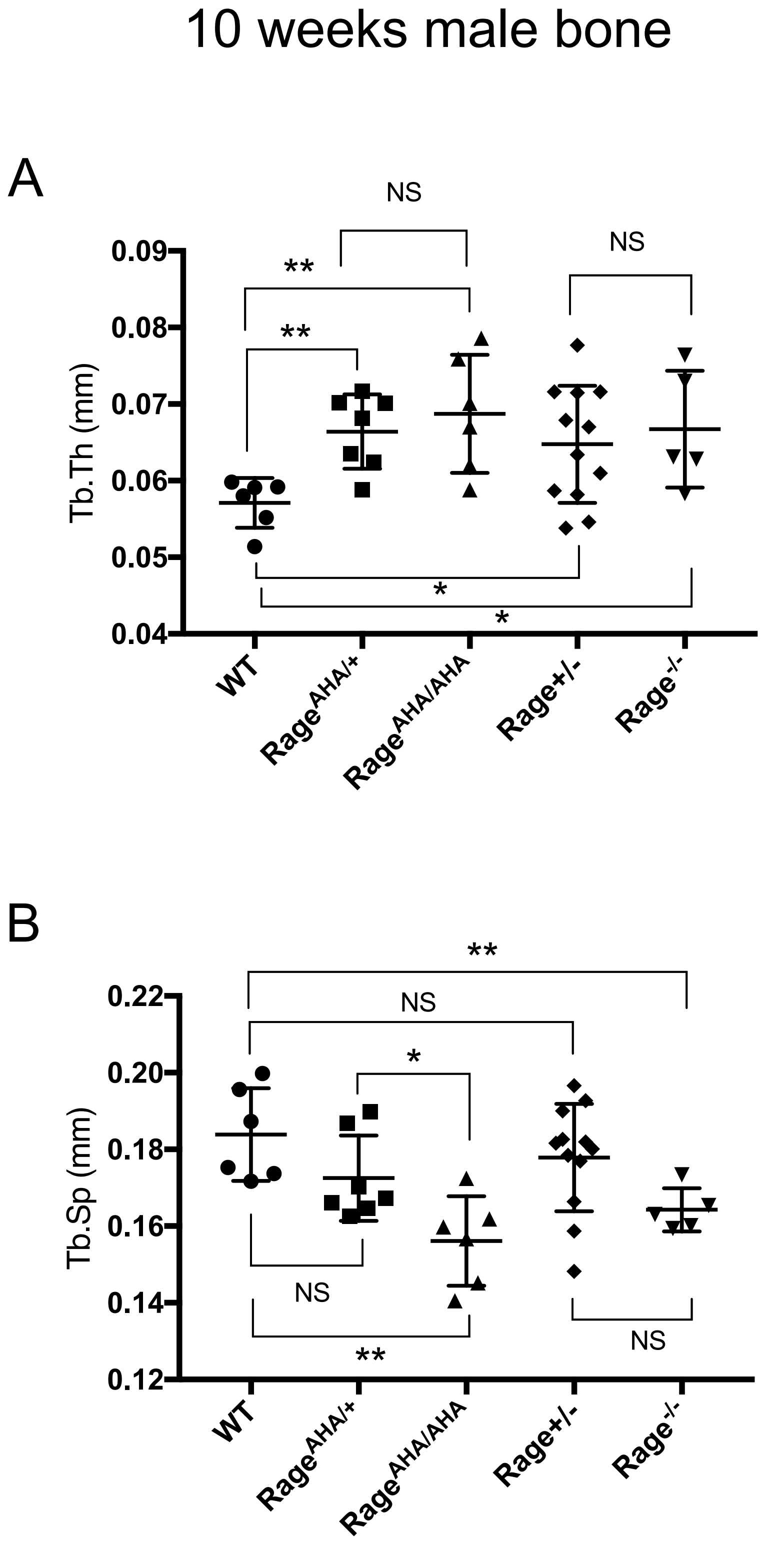


**Supplemental Figure S1. Trabecular bone morphometric analysis of 10-week old male WT, *Rage*$^{AHA/AHA}$, *Rage*$^{AHA/+}$, *Rage*$^{-/-}$ and *Rage*$^{+/-}$ mice.** (A) Trabecular thickness (Tb.Th), and (B) Trabecular separation (Tb.sp) as determined by $\mu$CT analysis. Error bars represent SD. *, ** and *** represents P < 0.05, 0.01 and 0.001 respectively.

Supplemental Figure S2

10 weeks female bone

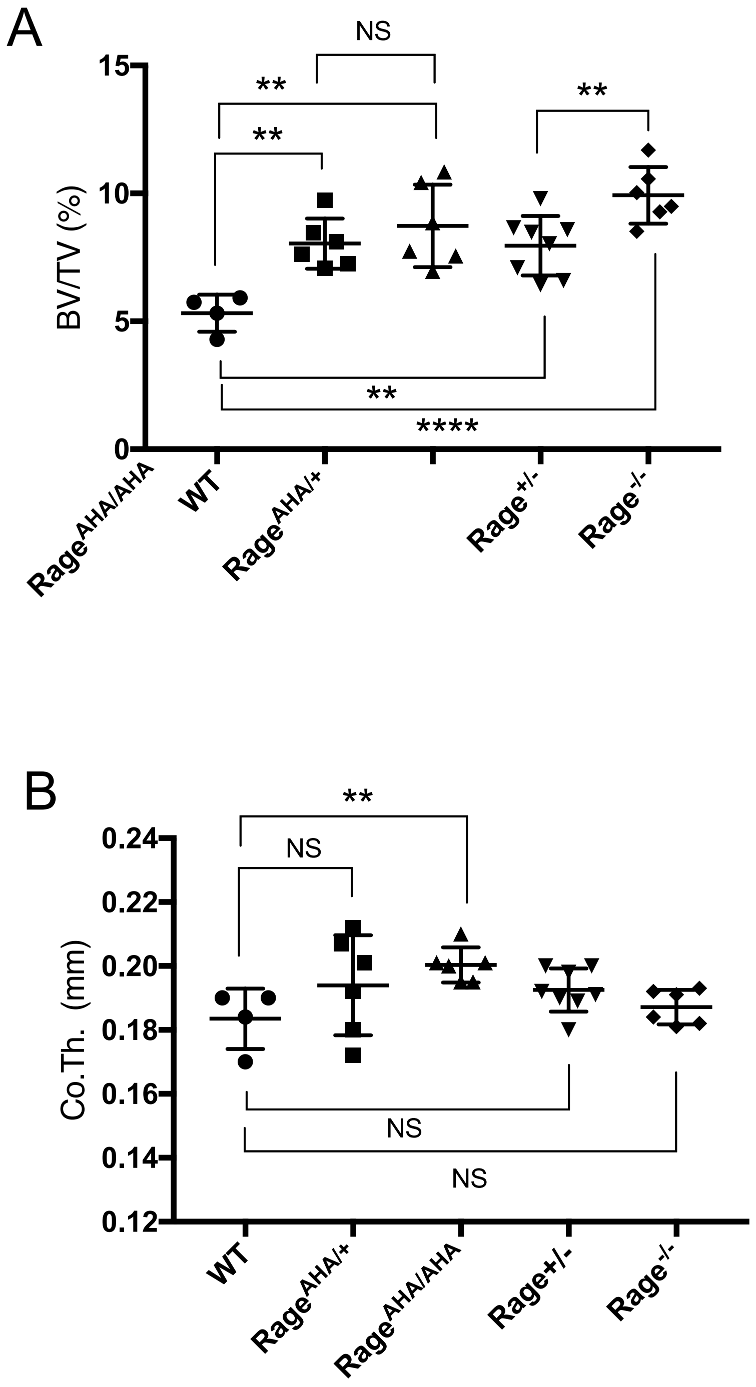


**Supplemental Figure S2. Female *Rage*$^{AHA/AHA}$ and *Rage*$^{AHA/+}$ mice develop osteopetrotic phenotype**. (A) Trabecular bone volume / tissue volume ratio (BV/TV), and (B) cortical bone thickness of femurs from 10-week old female WT, *Rage*$^{AHA/AHA}$, *Rage*$^{AHA/+}$, *Rage*$^{+/-}$ and *Rage*$^{-/-}$ mice.

Supplemental Figure S3

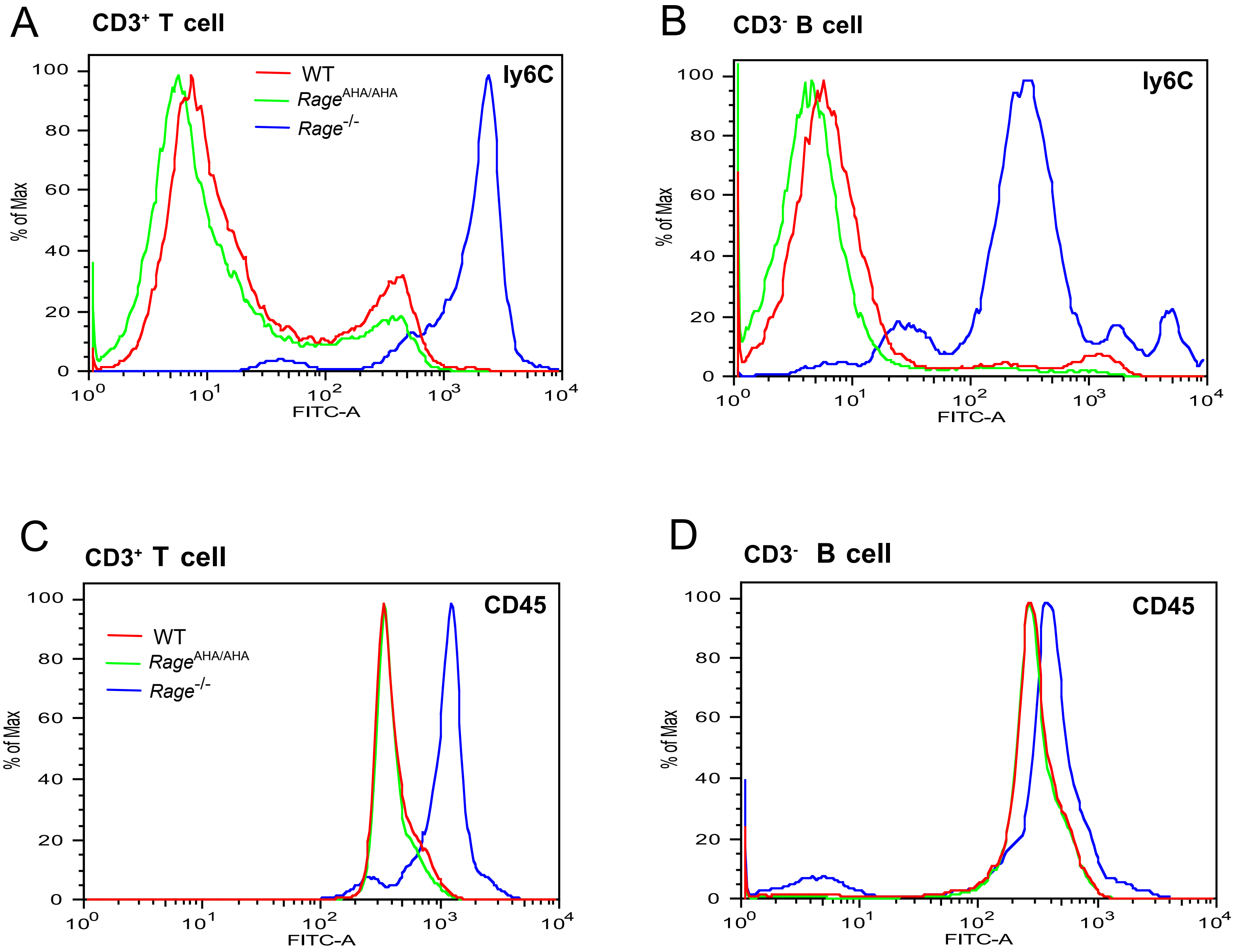


**Supplemental Figure S3. T cells and B cells from *Rage*$^{-/-}$ mice display greatly elevated surface expression of Ly6C and CD45.** (A) CD3+ T cells and (B) CD3- B cells from WT, *Rage*$^{AHA/AHA}$ and *Rage*$^{-/-}$ mice were stained with a rat anti-Ly6C-FITC mAb. (C) CD3+ T cells and (D) CD3- B cells were stained with a rat anti-CD45-FITC mAb. Cells isolated from 4 mice in each genotype were analyzed with almost identity result.

### Supplemental Figure S4

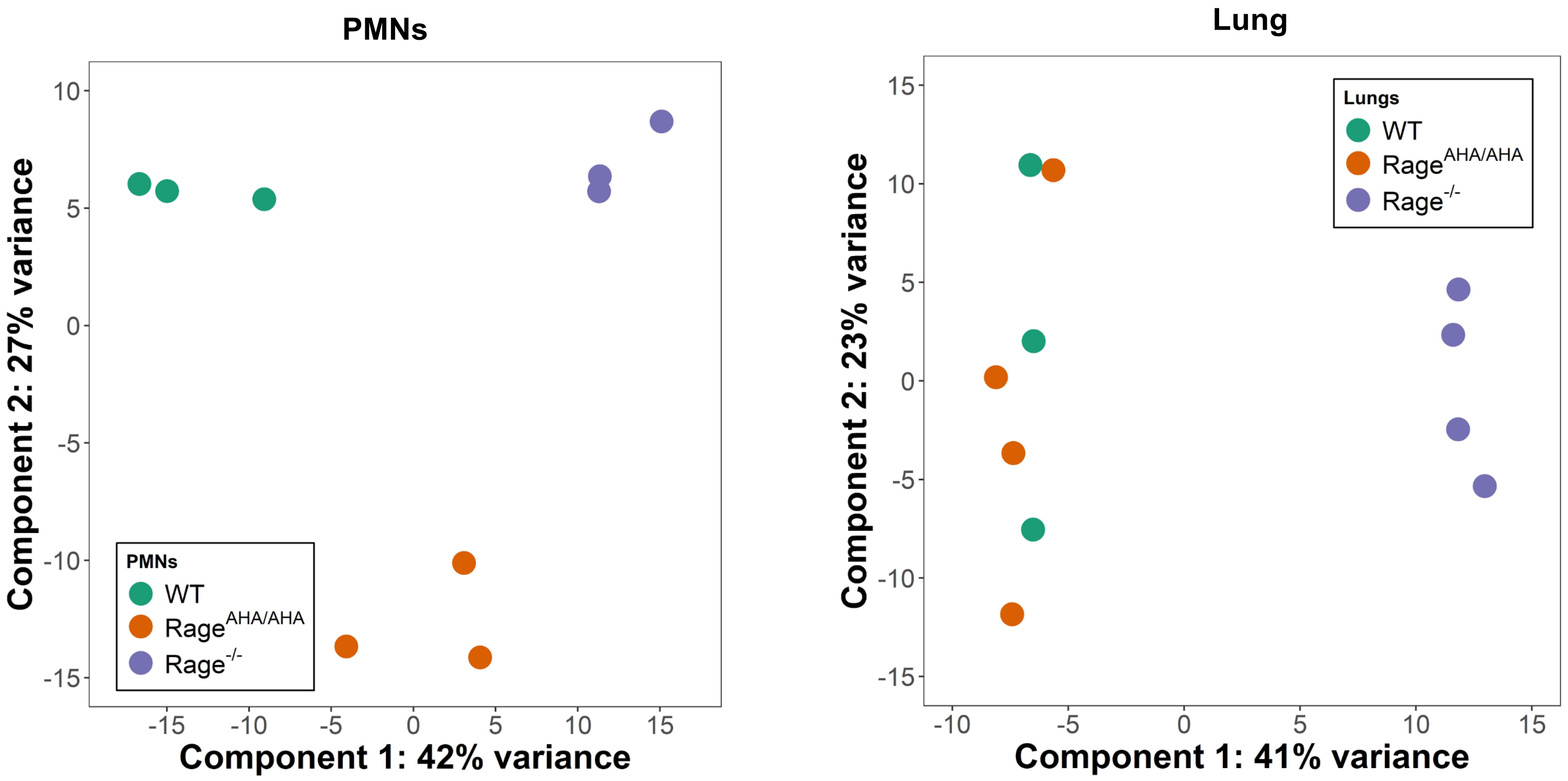


**Supplemental Figure S4. Principal component analysis of RNAseq data.** PMN and lung transcriptome of WT, $Rage^{AHA/AHA}$ and $Rage^{-/-}$ mice (3- 4 mice for each) was analyzed by principal component analysis using DESeq2 R package (v1.30.1). Delineation of the PMN transcriptome suggests clear separation between $Rage^{AHA/AHA}$ and $Rage^{-/-}$ compared to WT mice while the lung transcriptome suggest overlap between $Rage^{AHA/AHA}$ and WT mice with $Rage^{-/-}$ having clear separation.

### Supplemental Figure S5

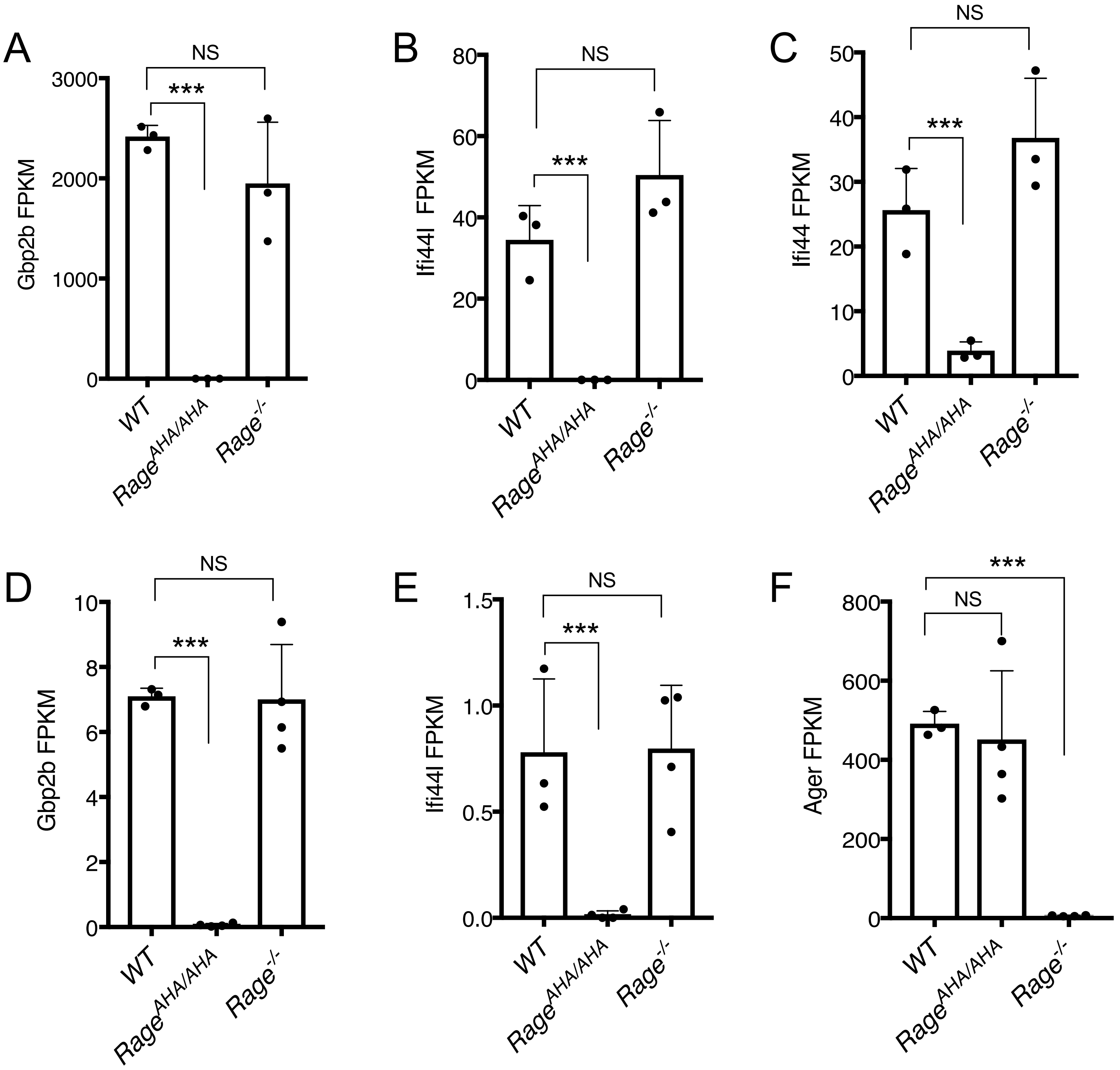


**Supplemental Figure S5. Expression levels of selected genes as determined by RNA-seq**. (A-C) Expression levels of Gbp2b, Ifi44L and Ifi44 in neutrophils, respectively, were plotted in FPKM (expected number of Fragments Per Kilobase of transcript sequence per Millions base pairs sequence). (D-F) Expression levels of Gbp2b, Ifi44L and Ager in lungs, respectively, were plotted in FPKM.
